# Rab14/MACF2/CAMSAP3 Complex Regulates Endosomal Targeting to the Abscission Site During Cytokinesis

**DOI:** 10.1101/2020.04.21.052449

**Authors:** Paulius Gibieža, Eric Peterman, Huxley K. Hoffman, Schuyler Van Engeleburg, Vytenis Arvydas Skeberdis, Rytis Prekeris

## Abstract

Abscission is complex cellular process that is required for mitotic division. It is well-established that coordinated and localized changes in actin and microtubule dynamics are vital for cytokinetic ring formation, as well as establishment of the abscission site. Actin cytoskeleton reorganization during abscission would not be possible without the interplay between Rab11- and Rab35-containing endosomes and their effector proteins, whose roles in regulating endocytic pathways at the cleavage furrow have now been studied extensively. Here, we identified Rab14 as novel regulator of abscission. We demonstrate that depletion of Rab14 causes either cytokinesis failure or significantly prolongs division time. We show that Rab14 regulates the efficiency of recruiting Rab11-endosomes to the central spindle microtubules and that Rab14 knockout leads to inhibition of actin clearance at the abscission site. Finally, we demonstrate that Rab14 binds to microtubule minus-end interacting MACF2/CAMSAP3 complex and that this binding is required for targeting of early endosomes to the central spindle. Collectively, our data identified Rab14/MACF2/CAMSAP3 as a protein complex that regulates Rab11-endosome targeting and the establishment of the abscission site.

## INTRODUCTION

Cytokinesis is the last stage of the cell cycle that leads to a physical separation of two daughter cells. Cytokinesis is initiated by a formation of cytokinetic actomyosin ring and contraction of this cytokinetic ring leaves two daughter cells connected with a thin intracellular bridge (ICB) that contains a microtubule-rich structure known as the midbody (MB) (Dionne *et al*, 2015). The resolution of the ICB, the process known as abscission, is the last step of cell division. It is becoming well-established that abscission depends on coordinated changes in cellular cytoskeleton, such as spastin-dependent cutting of central spindle microtubules and depolymerisation of actin cytoskeleton (Connell *et al*, 2009; Fremont *et al*, 2017; Guizetti *et al*, 2011; Prekeris 2011; Wilson *et al*, 2005). These localized changes in actin and microtubule cytoskeleton are vital to defining the abscission site and to regulating the timing of completing cell division (Addi *et al*, 2018; Fremont *et al*, 2017; Schiel and Prekeris 2010).

While it is now well-established that Rab11- and Rab35-containing endosomes are the key regulators of abscission (Prekeris 2011; Schiel and Prekeris 2013), many questions remain. For example, it remains to be fully understood how these endosomes are targeted specifically to the ICB. Indeed, Rab11 and Fip3 (Rab11-effector protein) were shown to be associated with specialized sub-population of recycling endosomes that are delivered to the abscission site only at late telophase (Schiel *et al*, 2012). Interestingly, during metaphase and anaphase Rab11/Fip3 associates with centrosomes (Collins *et al*, 2012), presumably preventing premature delivery of Rab11/Fip3-endosomes to the ICB. Furthermore, during early telophase, Rab11/Fip3-containing endosomes leave centrosomes and translocate to minus-ends of central spindle microtubules (Schiel *et al*, 2012; Simon *et al*, 2008). Although the functional consequence of this translocation remains to be fully defined, it is tempting to hypothesize that this recruitment of endosomes to microtubule minus-ends is required for efficient delivery of Rab11/Fip3 endosomes to the ICB and the midbody during abscission.

Since the roles of Rab11 and Rab35 in the control of cell division are well-defined, in this study we focus on identifying the possible functions of other Rabs in regulating the endosomal traffic to the abscission site during mitotic cell division. Recently we completed a proteomic analysis of post-abscission midbodies (MBs) and have shown that post-abscission MBs contain multiple endocytic Rabs (Peterman *et al*, 2019). In this study we test all of these MB-associated endocytic Rabs for their requirement during cytokinesis and abscission. We show that, in addition to Rab11 and Rab35, Rab14 also is required for abscission. We demonstrate that overexpression of Rab14 dominant-negative mutants, as well as knock-down (KD) or knock-out (KO) of Rab14 causes multi-nucleation and increase in time required for cells to divide. Also, we show that Rab14 functions upstream of Rab11, and that Rab14 also appears to mediate cytokinesis via actin clearance from the abscission sites. Finally, using co-immunoprecipitation/proteomic analysis, we identified the microtubule-bundling MACF2/CAMSAP3 protein complex as a Rab14 effector complex. Importantly, MACF2 knockdown leads to cytokinetic defects and inhibits the recruitment of Rab11/Fip3 endosomes to the ICB. Thus, we propose that Rab14/MACF2/CAMSAP3 complex mediates Rab11/Fip3-endosome targeting to central spindle microtubules within ICB and consequently regulates spatiotemporal regulation of abscission.

## RESULTS

### Rab14 is required for abscission

It is now well-established that endocytic transport plays an important role in regulating abscission; specifically, endocytic Rab11a/b and Rab35 were implicated to mediate this process (Fremont and Echard 2018; Gibieza and Prekeris 2018; Schiel *et al*, 2013). Since numerous Rab GTPases are known to regulate endocytic transport, we set out to identify other endocytic Rab GTPases that may contribute to mediating abscission. Recently, we completed proteomic analysis of post-abscission MBs purified from HeLa cells (Peterman *et al*, 2019), and so we first asked which Rab GTPases are present at the MBs and could play a role in abscission. In total, we identified 11 endocytic MB-associated Rabs GTPases (Figure 1A) (Peterman *et al*, 2019). Importantly, we identified Rab11b and Rab35, both known regulators of abscission (Collins *et al*, 2012; Dambournet *et al*, 2011; Fielding *et al*, 2005; Kouranti *et al*, 2006), as well as Rab8, which has also been shown to be present in the ICB (Schiel *et al*, 2012). Since Rab10, Rab14, Rab15, Rab21 and Rab29 have not been investigated for their involvement in regulating cell division, we next tested whether these Rabs may be required for completing cytokinesis. We have decided not to include Rab5 and Rab7 in this analysis since these two Rab GTPases regulate early endosome and lysosomal targeting, thus, their depletion could lead to wide-ranging defects and indirect effect on the abscission.

**Figure 1.**
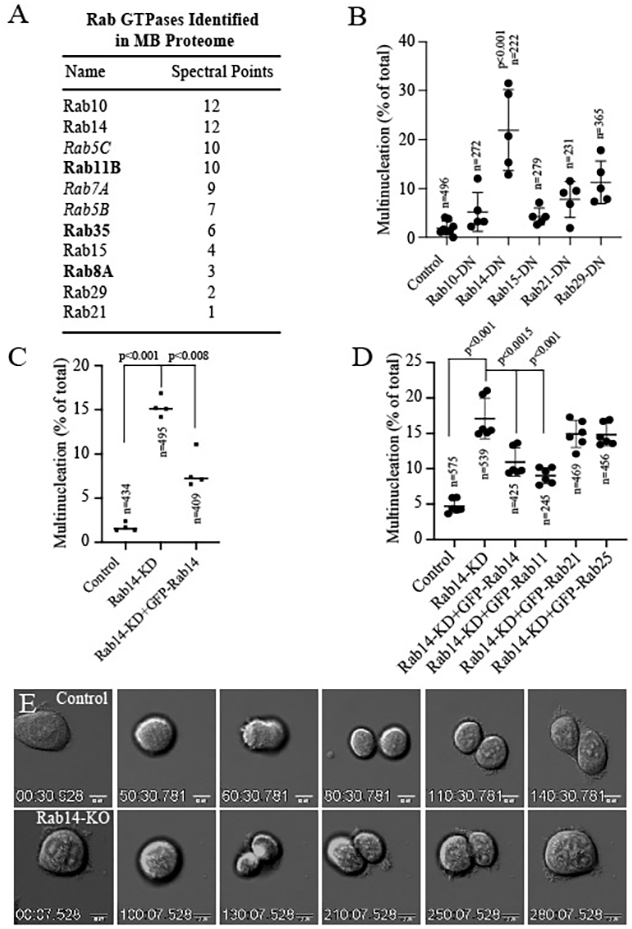
Rab14 is required for completion of cytokinesis. (A) Endocytic Rab GTPases present in post-mitotic MB proteome. (B) HeLa cells were transfected with dominant-negative mutants of various MB-associated Rabs. Multi-nucleated cells were then counted and expressed as percentage of total number of cells. Data shown are the means and standard deviations derived from three independent experiments. n is total number of cells counted. (C-D) Control and Rab14-KD cells were transfected with various GFP-tagged Rabs. The ability of overexpression to decrease multi-nucleation was then analysed. Data shown are the means and standard deviations derived from three independent experiments. n is total number of cells counted. (E) Individual images taken from time-lapse analysis of dividing control (top row) and Rab14-KO (bottom row) cells (also see Movies 1&2).

All Rab GTPases in cells cycle between GTP-bound (active), and GDP-bound (inactive) states. Thus, to test the involvement of aforementioned Rab GTPases in cytokinesis we created dominant-negative Rab10, Rab14, Rab15, Rab21 and Rab29 mutants by locking them in a GDP-bound state (S17N mutation) and performing a multi-nucleation assay. This assay is widely used as an indicator of defects in cytokinesis and we found out that amongst all the Rabs tested, only Rab14 dominant-negative mutant caused an increase in multi-nucleated cells (Figure 1B). Importantly, Rab14 knock-down (Supplemental Figure 1A) also led to increased multi-nucleation. The Rab14-KD induced cytokinetic defect could be rescued by over-expressing shRNA-resistant GFP-Rab14, but not GFP-Rab21 or GFP-Rab25 (Figure 1C-D).

While a multi-nucleation assay is a good initial assay to test for cytokinetic defects, it does not provide the information of what stage of cytokinesis is affected. If the defect is in abscission rather than cytokinetic ring formation and contraction, then dividing cells would be expected to accumulate in telophase. Consistent with Rab14 mediating the abscission, we found that Rab14-KD leads to increase in telophase cells (Figure 2B). Importantly, the abscission arrest induced by Rab14-KD was similar to the abscission arrest induced by Rab11a/b co-KD (Figure 2B), a well-established regulator of abscission (Schiel *et al*, 2013; Schiel and Prekeris 2010; Simon and Prekeris 2008; Wilson *et al*, 2005).

**Figure 2.**
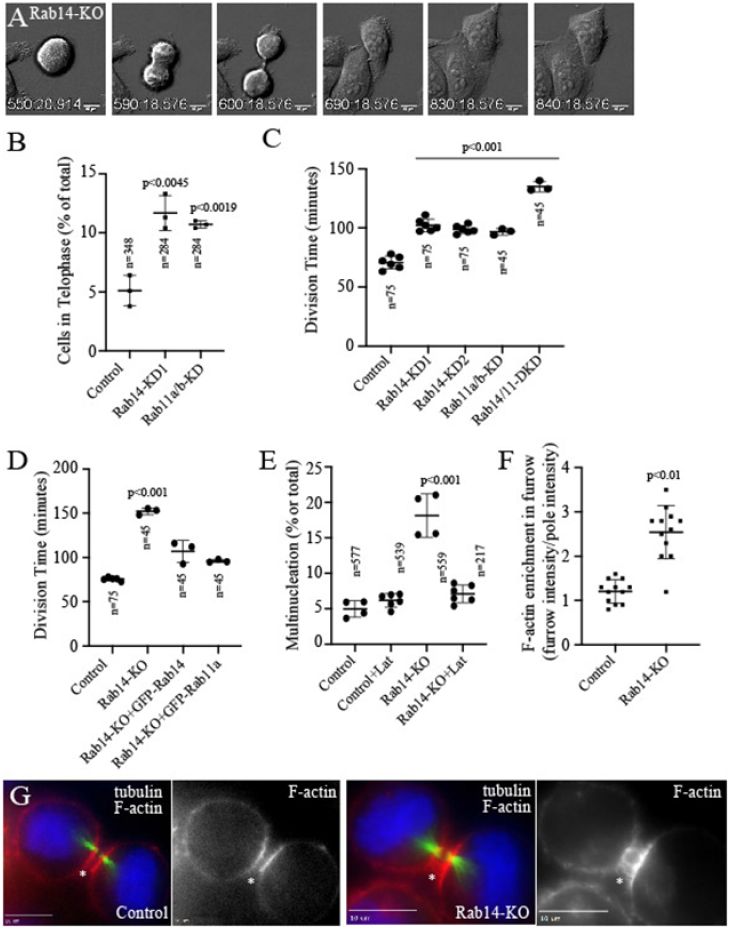
Rab14 knock-down increases time required for cells to undergo abscission. (A) Individual images taken from time-lapse analysis of dividing Rab14-KO cell (also see Supplemental Movie 3). (B) Control, Rab14-KD and Rab11a/b-KD cells were fixed and stained with anti-tubulin antibodies to identify cells in telophase. Data shown are the means and standard deviations derived from three independent experiments. n is total number of cells counted. (C-D) Control, Rab14-KD, Rab11a/b-KD, Rab14/11-DKD and Rab14-KO cells were analysed by time-lapse microscopy to measure the time needed to complete division (starting from metaphase). Where indicated, cells were transfected with GFP-Rab11a or GFP-Rab14. Data shown are the means and standard deviations derived from three independent experiments. n is total number of cells counted. (E) Control or Rab14-KO cells were incubated in the absence or presence of 4 nM Latrunculin A and multi-nucleated cells counted to assess their ability to complete cytokinesis. Data shown are the means and standard deviations derived from three independent experiments. n is total number of cells counted. (F-G) Control or Rab14-KO HeLa cells were fixed and stained with phalloidin-596 and anti-tubulin antibodies. The fluorescence of phalloidin-596 at the ICB was then evaluated. The quantification is shown at panel (F). Data shown are the means and standard deviations derived from three independent experiments.

To further understand the role of Rab14 in cytokinesis we next created two different HeLa Rab14 knock-out cell lines (Rab14-KO1 and Rab14-KO2, Supplemental Figure 1B-C) and performed a time-lapse analysis of cells during mitotic cell division. Consistent with an increase in multi-nucleation, about ~17% of Rab14-KO cells failed cytokinesis and regressed furrow to form bi-nucleate cells (Figure 1E; Movie 1&2). The remaining 83% of cells did eventually divide, but spent a much longer time in telophase (Figure 1E and Figure 2A, D; Movie 3). Importantly, similar increases in division time were also observed in Rab14-KD cells (Figure 2C) Abscission defects caused byRab14-KO could be rescued by over-expression of GFP-Rab14 (Figure 2D).

### Rab14 and Rab11 are part of the same abscission regulatory pathway

Previous studies have demonstrated that Rab11-endosomes translocate from centrosomes to the ICB and the midbody to mediate actin clearance and the establishment of the abscission site (Schiel *et al*, 2012; Simon *et al*, 2008). Thus, we next decided to investigate the possible interplay between Rab14 and Rab11 function during abscission. Our data demonstrate that both Rab11-KD and Rab14-KD lead to delays in division time (Figure 2C), and so we next tested the effect of treating HeLa-Rab14-KD cells with Rab11a/b siRNA. As shown in Figure 2C, triple KD of Rab11a, Rab11b and Rab14 further increased the time required for cell division, indicating that Rab14 and Rab11 may function in parallel or the same pathway.

To further investigate interplay between Rab14 and Rab11, we overexpressed GFP-Rab11a in either Rab14-KD or Rab14-KO cells and analysed division time, as well as multi-nucleation. As shown in Figures 1D and 2D, overexpression of GFP-Rab11a was able to rescue cytokinetic defects induced by Rab14 depletion.

Previous studies have shown that Rab11a/b play an important role in cell division by regulating actin clearance from the ICB during cytokinesis (Dambournet *et al*, 2011; Fielding *et al*, 2005; Wilson *et al*, 2005). In order to find out if Rab14 is also required for actin clearance at the abscission site we co-stained control or Rab14-KO cells with anti-tubulin antibodies and phalloidin-Alexa596 (F-actin marker). Consistent with the hypothesis that Rab11 and Rab14 may be acting in the same pathway, Rab14 depletion led to an increase in F-actin in the ICB (Figure 2F-G). Finally, to examine whether the aberrant F-actin accumulation was the cause of the Rab14 depletion induced cytokinesis defects, we treated control and Rab14-KO cells with low dose (4 nM) of the F-actin depolymerizing drug LatrunculinA (LatA), and then tested the ability of cells to complete cytokinesis by multi-nucleation assay. As shown in Figure 2E, LatA treatment rescued Rab14-KO induced multi-nucleation while having no effect on multi-nucleation in control cells.

### Rab14 localizes to early endosomes that translocate from centrosomes to minus-ends of central spindle microtubules during telophase

Our data so far indicate that Rab14 is required for successful completion of abscission and that, just like Rab11, Rab14 appears to function by regulating actin disassembly at the abscission site. The question that does remain is whether Rab14 mediates a distinct pathway of regulating actin dynamics that is parallel to Rab11-dependent pathway or if Rab14 and Rab11 are steps in same endocytic cascade that eventually leads to actin clearance at the abscission site. To identify Rab14-dependent cytokinetic steps, we first analysed Rab14 localization during cell division. To that end, we used CRISPR/Cas9 to create a HeLa cell line with GFP-tagged endogenous Rab14 (Supplemental Figure 2B). We then followed GFP-Rab14 localization at the different stages of mitotic cell division. As shown in Figure 3A, Rab14 is mostly cytosolic (presumably GDP-bound and inactive) during metaphase and anaphase. During telophase Rab14 accumulates around centrosomes, as well as on endosome-like structures located in close proximity of minus-ends of central spindle microtubules (Figure 3A). Interestingly, it has been shown by several studies that Rab11-containing recycling endosomes also accumulate at minus-end of microtubules and that this type of accumulation may be required for efficient delivery of recycling endosomes to the ICB and the MB (Schiel *et al*, 2012; Simon *et al*, 2008). To determine the identity of these Rab14-containing organelles we next co-stained telophase and interphase cells with EEA1 (early endosome marker). As shown in Figure 3B, GFP-Rab14 almost completely colocalized with EEA1, demonstrating that Rab14 is located at the early endosomes and that these early endosomes are targeted to the close proximity of minus-ends of the central spindle during telophase. Similarly, in interphase cells, Rab14 also colocalized with EEA1, further supporting the findings that Rab14 is predominately early endosomal protein (Supplemental Figure 2A).

**Figure 3.**
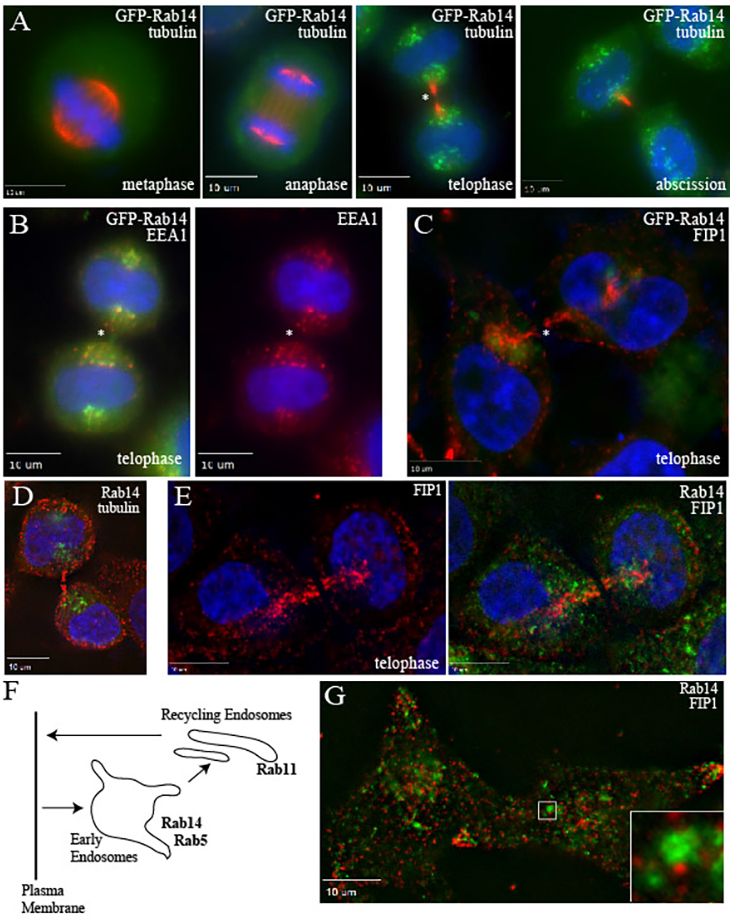
Rab14 is enriched at early endosomes located at the minus-ends of central spindle microtubules. (A-C) HeLa cells expressing GFP-tagged endogenous Rab14 were fixed and stained with anti-tubulin (A), anti-EEA1 (B) or anti-Fip1 (C) antibodies. Asterisks mark the ICB. (D-E) HeLa cells in telophase were fixed and stained with anti-Rab14 (D and E), anti-tubulin (D) or anti-Fip1 (E) antibodies. (E) Schematic representation of proposed endocytic Rab14 localization. (F) HeLa cells in interphase were fixed and stained with anti-Fip1 and anti-Rab14 antibodies.

To dissect the possible connection between Rab11 and Rab14 endosomes we next co-stained cells with anti-Fip1 antibodies. Fip1 is a well-established Rab11-effector that is present on most Rab11-containing recycling endosomes (Junutula *et al*, 2004). While Fip1 does not appear to regulate abscission (Wilson *et al*, 2005), it can be used as a marker for Rab11-containing recycling endosomes during interphase and mitotic cell division (Peden *et al*, 2004). Consistent with our data described above, we observed little colocalization between GFP-Rab14 and Fip1 during telophase (Figure 3C). Importantly, similar results were observed when cells were stained with anti-Rab14 antibody (Figure 3D-G). While we observed little colocalization between Rab14 and Fip1, we could observe localization of Rab14 and Fip1 endosomes in close proximity, especially at the minus-ends of central spindle microtubules (Figure 3C and E). We often observed what appears to be a Fip1-endosome budding from Rab14-containing early endosome (Figure 3G, see inset). These observations are fully consistent with the well-established fact that Rab11-recycling endosomes bud from Rab5 (and presumably Rab14) containing early endosomes (Figure 3F).

### Rab14 is required for targeting of Rab11/Fip3 endosomes at the minus-ends of central spindle microtubules

Our localization data suggests that Rab11-recycling endosomes bud directly from Rab14-containing early endosomes that are situated at the minus-ends of central spindle microtubules during telophase. This raises an interesting possibility that positioning Rab14-endosomes at the minus ends of the central spindle may regulate the targeting of newly formed Rab11-recycling endosomes to the ICB and the midbody. To test this hypothesis, we investigated the localization of Rab11/Fip1-endosomes during telophase in control and Rab14-KO cells. Since some reports suggested that Rab14 can directly bind to Fip1 (Kelly *et al*, 2009; Lall *et al*, 2015), we first tested whether Rab14 KO affects Fip1 recruitment to endosomes. As shown in Supplemental Figure 2C, Fip1 targeting to endosomes in interphase cells was independent of Rab14. Next, we analysed the distribution of Fip1 during telophase. As shown in Figure 4A-B, in agreement with previously published reports, Rab11-recycling endosomes (as determined by Fip1 localization) accumulate at the central spindle during telophase and this accumulation is inhibited by Rab14 KO.

**Figure 4.**
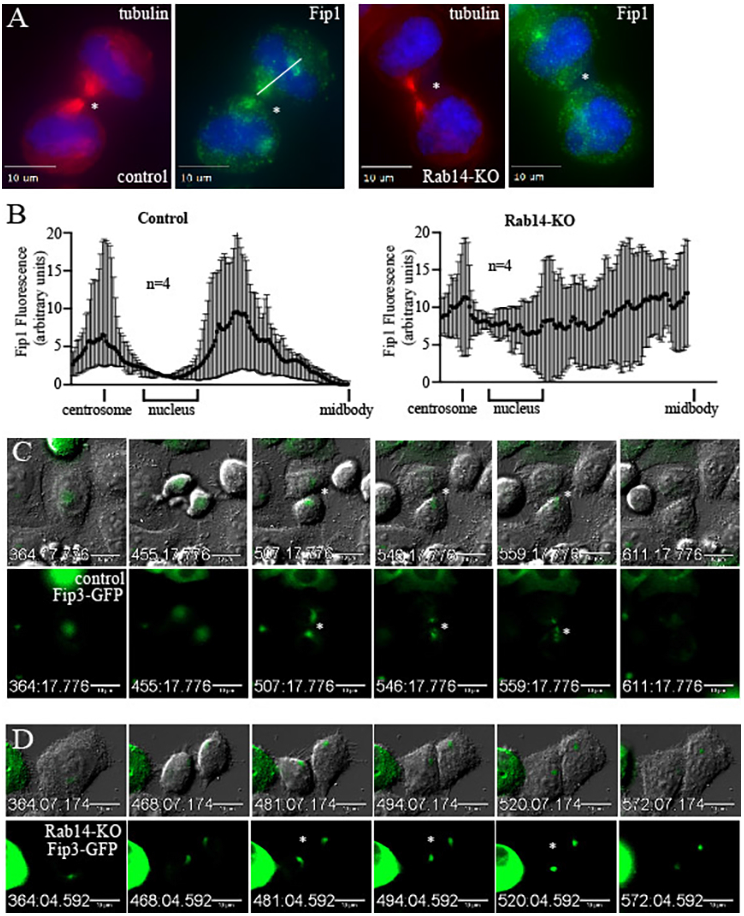
Rab14 is required for targeting Rab11-endosomes to the central spindle microtubules. (A-B) Control or Rab14-KO HeLa cells were fixed and stained with anti-tubulin and anti-Fip1 antibodies. Line-scan analysis of Fip1 fluorescence was then performed in telophase cells (see line in panel A). Asterisks mark the ICB. Data shown in (B) are the means and standard deviations derived from line-scans of four different cells. (C-D) Time-lapse images of dividing control or Rab14-KO cells expressing GFP-Fip3. Asterisks mark the ICB.

To further investigate the effect of Rab14 KO on recycling endosome targeting to the ICB we transfected cells with GFP-Fip3 and analysed recycling endosome dynamics by time-lapse microscopy. Fip3 is also a Rab11-interacting protein that has been directly implicated in regulating abscission by delivering p50RhoGAP and contributing to actin depolymerisation at the abscission site (Schiel *et al*, 2012). Unlike Fip1 that localizes to most Rab11-recycling endosomes, Fip3 only localizes to a subset of recycling endosomes that are targeted to the abscission site (Prekeris 2015; Simon *et al*, 2008), thus, Fip3 is an excellent marker to monitor the dynamics of these endosomes during cytokinesis. As shown in Figure 4C (also see Movies 4&5), in control HeLa cells GFP-Fip3-endosomes move from centrosomes to the ICB as cell completes division (in 81.5% of cells; 27 dividing cells analysed). In contrast, in Rab14-KO cells, GFP-Fip3-endosomes accumulate at the centrosome, but then fail to re-localize to the ICB at late telophase (in 90% of cells; 20 dividing cells analysed) (Figure 4D; Movies 6&7). Taken together, our data suggests that Rab14 is required for targeting of early endosomes to the minus-ends of central spindle microtubules and consequently affecting the delivery of Rab11/Fip3 endosomes to the abscission site.

### MACF2 is Rab14-binding protein that is required for abscission

While our data demonstrate that Rab14 is required for endosome targeting to minus-ends of central spindle microtubules, the molecular machinery governing Rab14 function during telophase remains unclear. Keeping in mind that all Rab proteins function via GTP-dependent interactions with a diverse array of specific effector proteins (Hutagalung and Novick 2011), we decided to identify Rab14 effector proteins that function in cell division. To that end, we next created HeLa cells stably expressing either FLAG-Rab14 or FLAG-Rab14-Q70L (constitutively active Rab14 mutant) and performed co-immunoprecipitation/proteomics analysis from synchronized HeLa cells in telophase using anti-FLAG antibody. Since Rab GTPases bind to their effectors in GTP-dependent manner we filtered all candidate proteins based on two main criteria: (1) the candidate proteins should not be present at IgG control and (2) candidate proteins should be enriched at least 2X in the Rab14-Q70L sample (Figure 5A). Proteomic analysis identified 9 candidates that appear to be interacting specifically with GTP-bound Rab14 (Figure 5A). One of these proteins, MACF2 (microtubule actin cross-linking factor 2), is a large scaffolding protein that belongs to a spectraplakin family and is implicated in microtubule bundling, cross-talk between actin and microtubules, as well as regulating non-centrosomal microtubule dynamics (Noordstra *et al*, 2016; Sun *et al*, 2001). Furthermore, MACF1 (also known as ACF7) is a closely related MACF2 isoform that was shown to bind to CAMSAP3/Patronin proteins and interact with microtubule minus-ends (Noordstra *et al*, 2016). Finally, MACF1 was shown to be required for Rab11-endosome traffic during epithelia polarization (Noordstra *et al*, 2016). This raises an interesting possibility that MACF2 may regulate Rab11-endosome targeting during cytokinesis. Testing this hypothesis is the focus of the rest of this study.

**Figure 5.**
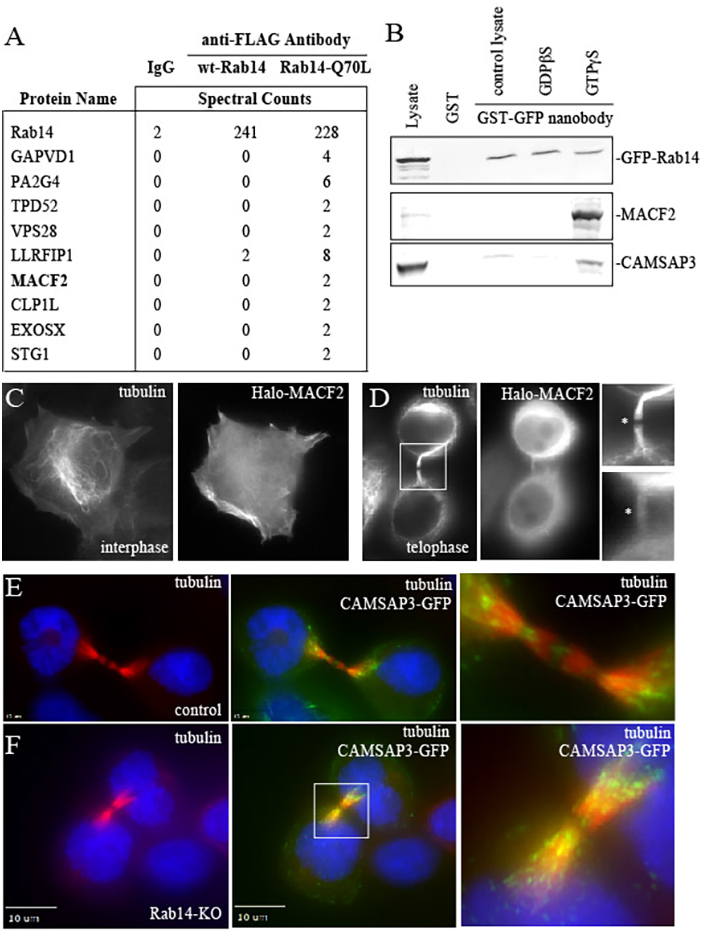
Rab14 binds to MACF2/CAMSAP3 complex. (A) Putative Rab14-interacting proteins identified in FLAG-Rab14 immunoprecipitation and proteomic analysis. (B) Lysates from HeLa cells expressing endogenously labelled GFP-Rab14 were incubated with either glutathione beads coated with GST or GST-anti-GFP-nanobody. The beads were then washed and bound protein analysed by western blotting. (C-D) HeLa cells transfected with Halo-MACF2 were fixed and stained with Halo ligand and anti-tubulin antibodies. (E-F) Control (E) or Rab14-KO (F) HeLa cells were transfected with GFP-CAMSAP3, then fixed and stained with anti-tubulin antibodies.

To confirm that MACF2 is a Rab14 interacting protein, we used a GST-tagged anti-GFP nanobody to pull-down endogenously tagged GFP-Rab14 using glutathione bead pull-down assay, followed by immunobloting using anti-MACF2 antibodies. As shown in Figure 5B, MACF2 was pulled-down only when Rab14 was loaded with non-hydrolysable GTP analogue, GTPyS, an observation consistent with MACF2 being a Rab14 effector protein. Since MACF1 was shown to interact with CAMSAP3, a microtubule minus-end interacting protein, we wondered whether MACF2 and Rab14 complex also interacts with CAMSAP3. Consistent with this hypothesis, anti-GFP-nanobodies co-immunoprecipitated CAMSAP3 with GFP-Rab14. Importantly, CAMSAP3 also co-precipitated predominately with GTP-locked GFP-Rab14 (Figure 5B), suggesting that MACF2 and CAMSAP3 interacts with activated Rab14, thus is Rab14 effector complex.

To determine whether MACF2 is important for cytokinesis, we next transfected cells with Halo-MACF2 or GFP-CAMSAP3 and analysed their localization during telophase in HeLa cells. As previously reported (Sun *et al*, 2001), MACF2 partially colocalized with tubulin in interphase cells (Figure 5C). Similarly, Halo-MACF2 could also be observed in the ICB during telophase (Figure 5D). Interestingly, GFP-CAMSAP3 was highly enriched at the central spindle microtubules (Figure 5E), consistent with its role in regulating/protecting minus-ends of non-centrosomal microtubule bundles (Noordstra *et al*, 2016). Importantly, Rab14 KO did not have any effect on Halo-MACF2 or GFP-CAMSAP3 localization (data not shown and Figure 5F) suggesting that both, MACF2 and CAMSAP3 are targeting to the ICB independent of Rab14 (presumably by direct binding to microtubules). Thus, we suggest that MACF2 and CAMSAP3 together play a role in modulating abscission.

In order to better understand MACF2 functions during cell division, we used two different shRNAs (MACF2-KD1 and MACF2-KD2) to deplete MACF2 in HeLa cells (Supplemental Figure 3). Once again, we performed time-lapse analysis to test the effect of MACF2 depletion on time required for cells to complete abscission. As shown in Figure 6A-B, (also see Supplemental Movie 8), MACF2 depletion increased overall cell division time from 70 min. in control cells, to 132 min. in HeLa-MACF2-KD1 and to 124 min. in HeLa-MACF2-KD2 cells. Overall, these results suggest that MACF2 is a Rab14 effector protein and is another important regulator of cell division.

**Figure 6.**
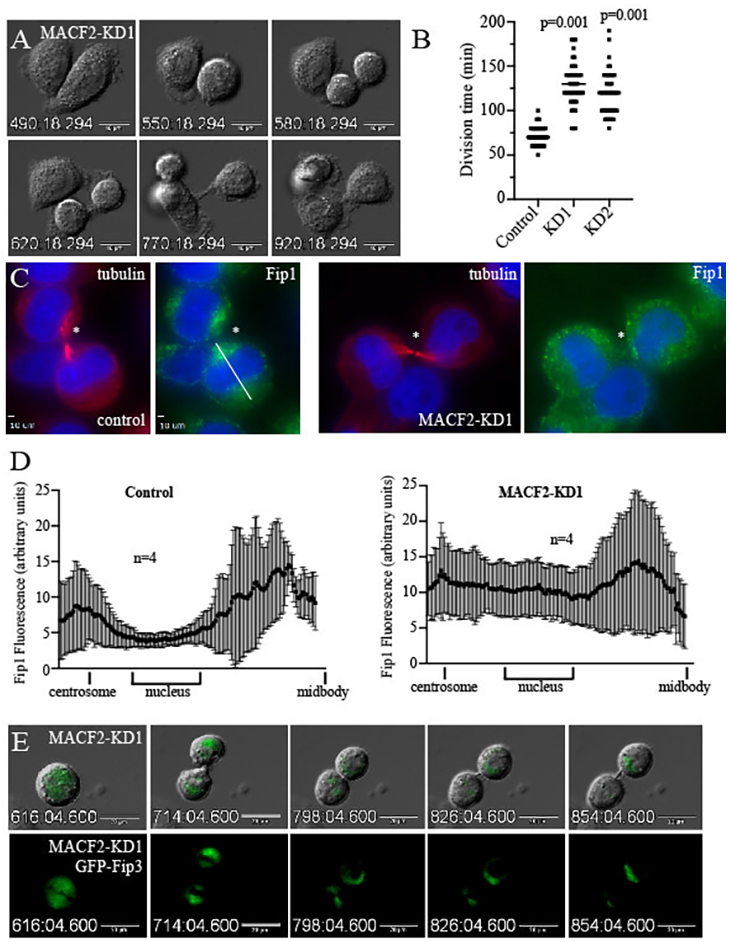
MACF2 is required for completion of cytokinesis and for targeting of Rab11-endosomes to the central spindle microtubules. (A-B) Time-lapse images of dividing MACF2-KD1 HeLa cell. Panel (B) shows quantification of time required for completing division (starting from metaphase). Data shown in (B) are the means and standard deviations derived from three independent experiments. (C-D) Control or MACF2-KD1 cells were fixed and stained with anti-tubulin and anti-Fip1 antibodies. Line-scan analysis of Fip1 fluorescence was then performed in telophase cells (see line in panel C). Asterisks mark the ICB. Data shown in (D) are the means and standard deviations derived from line-scans of four different cells. (E) Time-lapse images of dividing MACF2-KD1 cells expressing GFP-Fip3.

### MACF2 is required for Rab11/Fip3-endosome targeting to the ICB

Since Rab14 depletion does not affect MACF2 targeting to the ICB we hypothesized that MACF2 may function as a tether anchoring Rab14-containing early endosomes at the central spindle during early telophase. To test this hypothesis, we stained control or MACF2-KD cells with anti-Fip1 antibody to analyse the localization of Rab11-endosomes during telophase. Consistent with the involvement of MACF2 in regulating endocytic transport to the ICB, MACF2 depletion inhibited accumulation of Rab11/Fip1-endosomes at the minus-ends of central spindle microtubules (Figure 6C-D).

We next wondered whether the depletion of MACF2 would also affect trafficking of Rab11/Fip3-endosomes to the abscission sites at the ICB. For this purpose, we transfected control and MACF2-KD1 cells with GFP-FIP3 construct and performed time-lapse imaging analysis. As shown in the Figure 6E, FIP3 localisation at the ICB during late telophase was disrupted by the depletion of MACF2 (in 85.2% of cells; 27 dividing cells analysed) (also see Movie 9). Altogether, these results suggest that MACF2 and CAMSAP3 complex regulates Rab14 and Rab11-endosome targeting to the central spindle microtubules during early telophase, thus, mediating the actin clearance and establishment of the abscission site during mitotic cell division.

## DISCUSSION

Abscission is the last step of mitotic cell division that leads to resolution of the intracellular bridge (ICB) and physical separation of two daughter cells. It is now becoming clear that abscission is a very complex and highly regulated event (Addi *et al*, 2018; Fremont and Echard 2018; Schiel and Prekeris 2010). Indeed, recent studies identified a new mitotic checkpoint, known as abscission checkpoint (also sometimes referred to as NoCut), that is activated by lagging chromosomes and leads to the arrest of the cells in telophase (Bai *et al*, 2020; Petsalaki and Zachos 2020; Sadler *et al*, 2018). We now know that abscission involves highly organized remodelling of the cytoskeleton at the ICB. Specifically, the abscission site is defined by localized disassembly of actin cytoskeleton that is presumably a remnant of the cytokinetic contractile ring (Addi *et al*, 2018; Fremont and Echard 2018; Fremont *et al*, 2017; Schiel and Prekeris 2013; Schiel *et al*, 2012). Actin depolymerisation then leads to spastin-dependent microtubule severing and formation of ESCRT polymers that eventually drives plasma membrane fusion and resolution of the ICB (Schiel *et al*, 2013). Here we define a new Rab14 and MACF2/CAMSAP3-dependent pathway that is needed for F-actin clearance from the abscission site and for successful completion of cytokinesis.

Recycling endosomes have emerged as key regulators of localized actin disassembly and, consequently, completion of abscission. It was shown that various regulators of actin depolymerisation, such as MICAL1, p50RhoGAP and OCRL, are all delivered by two distinct sets of recycling endosomes, Rab11 and Rab35 endosomes (Collins *et al*, 2012; Dambournet *et al*, 2011; Fremont *et al*, 2017; Schiel *et al*, 2012). It is now believed that rapid and highly targeted delivery of these actin regulators to the abscission site is dependent on central spindle microtubules to transport of Rab11 and Rab35 endosomes rather than relying on passive diffusion from cytosol to the microtubule-packed ICB. Central spindle microtubules are perfectly suited for this task since they are arranged in polarized fashion with plus-ends oriented toward the center of the ICB where they form a unique structure known as the midbody (MB). What remains to be fully understood is how these endosomes are targeted to the central spindle microtubules where they can engage with Kinesin II molecular motors to be delivered to the forming abscission site. Interestingly, numerous studies have shown that during metaphase and anaphase Rab11-endosomes accumulate around the centrosome, although the machinery mediating centrosomal accumulation remains unclear (Figure 7) (Collins *et al*, 2012; Schiel *et al*, 2012). Once a cell enters telophase, Rab11 endosomes leave the centrosome and transiently accumulate around minus-ends of central spindle microtubules before entering the ICB and mediating cell abscission (Figure 7) (Hoog *et al*, 2011; Simon *et al*, 2008). The function of this endosome accumulation at the central spindle minus-ends remains unclear, but it is tempting to hypothesize that this accumulation helps with loading of Rab11-endosomes for plus-end directed (kinesin-dependent) transport to the abscission site. In this study we identify Rab14 as a protein required for endosome accumulation at the central spindle and also demonstrate that depletion of Rab14 leads to delay or failure of abscission.

**Figure 7.**
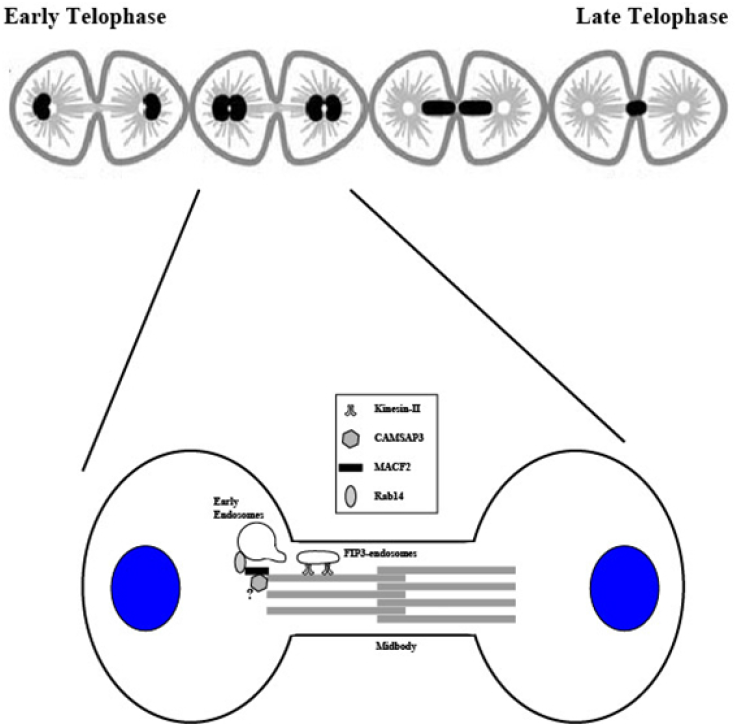
Schematic representation of the proposed function for Rab14/MACF2 complex. MACF2 is recruited to the central spindle microtubules either by directly binding to microtubule bundles or by co-binding to CAMSAP3, the protein that interacts with microtubule minus-ends. MACF2 then interacts with Rab14, thus, recruiting Rab14-containing early endosomes to the central spindle. Rab11/Fip3 recycling endosomes then buds from early endosomes and is delivered to the abscission site via kinesin-2 molecular motor.

We originally identified Rab14 as one of several endocytic Rabs present in the MB proteome (Figure 1A) (Peterman *et al*, 2019). Importantly, our analysis shows that, in addition to Rab11 and Rab35 (also identified in MB proteome), Rab14 is the only MB-associated Rab that is required for abscission, since depletion of Rab14 or expression of Rab14 dominant-negative mutant leads to a significant increase in time required for abscission or complete abscission failure. These data led us to hypothesize that Rab14 may also mark the endosomes (possibly even Rab11-endosomes) that may deliver yet another actin regulator to the abscission site. Surprisingly, our time-lapse and immunofluorescence analyses show that Rab14 is rarely observed inside the ICB, but instead accumulates at the minus-ends of the microtubules. Additionally, Rab14 appears to be predominantly present on early endosomes that also contain EEA1 and presumably Rab5. Since it is now well-established that Rab11-endosomes bud from early endosomes we hypothesized that Rab14 may mediate targeting of early endosomes to the minus-ends of central spindle microtubules, thus, ensuring efficient formation and targeting of Rab11-endosomes to the central spindle and the abscission site (Figure 7). Consistent with this hypothesis, Rab14 depletion decreased Rab11-endosome accumulation at the central spindle, as well as inhibited actin disassembly at the ICB during abscission. Furthermore, Rab14 KO-dependent abscission phenotypes can be rescued by over-expressing of Rab11, indicating that Rab11 functions downstream of Rab14.

To determine how Rab14 itself is targeted to minus-ends of microtubules we next completed Rab14 immunoprecipitation/proteomic analysis using HeLa cells synchronized in telophase. Importantly, proteomic analysis followed by pull-down assays identified MACF2 as a Rab14-ineracting protein. What makes MACF2 such an interesting protein is that is known to bind both, actin and microtubules, as well as mediate microtubule bundling. Additionally, MACF1, a closely related MACF2 paralogue, was shown to bind to CAMSAP3 and mediate targeting of MACF1/CAMSAP3 to minus-ends of microtubules in epithelial cells (Noordstra *et al*, 2016). Since CAMSAP3 co-immuno-precipitates with Rab14 and MACF2, we propose that MACF2/CAMSAP3 complex tethers Rab14 to the minus-ends of central spindle microtubules, mediating targeting of early endosomes and consequently Rab11-endosomes to the central spindle and abscission site (Figure 7). Consistent with this hypothesis, we show that CAMSAP3, and to a lesser degree MACF2, localize to the central spindle during abscission. Importantly, MACF2 depletion also inhibits Rab11-endosome accumulation at the central spindle and also leads to increase in time required to complete abscission.

Based on all our data we propose that Rab14, MACF2 and CAMSAP3 complex formation plays an important role in targeting endosomes to the central spindle, thus, contributing to efficient “loading” of Rab11-endosomes on kinesins and driving Rab11/Fip3-dependent delivery of p50RhoGAP to the abscission site. Many questions, however, remain and further studies will be needed to answer them. Does Rab14 also regulate targeting of Rab35 endosomes? What drives formation of Rab14/MACF2/CAMSAP3 complex during telophase, but blocks it from functioning in metaphase and anaphase? Interesting, Rab14 was also implicated in regulating epithelial cell polarity (Lu and Wilson 2016), as well as recycling of integrins during cell migration (Gundry *et al*, 2017). Both of these cellular processes are also dependent on actin and microtubule dynamics, thus, it will be interesting to see whether Rab14 interaction with MACF2/CAMSAP3 complex may have functions during cell polarization and migration.

## MATERIALS AND METHODS

### Cell culture and treatments

Cells were grown in 37 °C humidified incubator at 5% CO_2_, routinely tested for mycoplasma, and were maintained in DMEM with 10% FBS and 1% penicillin/streptomycin. To create FLAG-Rab14 stable cell line, HeLa cells were infected with lentivirus pLVX:FLAG-Rab14. To create FLAG-Rab14-Q70L stable cell line, HeLa cells were infected with lentivirus pLVX:FLAG-Rab14-Q70L. Stable Rab14 knockdown HeLa cell lines (HeLa-Rab14-KD1 and HeLa-Rab14-KD2) were generated by using Sigma lentiviral human Rab14 shRNA plasmids (TRCN0000293339 and TRCN000293336). The population was selected with puromycin, and stable clones were isolated. Stable Rab14 knockout HeLa cell line (HeLa-Rab14-KO) was generated by using Dharmacon human Rab14 crRNAs (51552). For rescue experiments, GFP-Rab14, GFP-Rab11a or GFP-Rab35 constructs were transiently transfected into HeLa-Rab14-KD2 or HeLa-Rab14-KO cells by using jetPRIME (Polyplus transfection kit). Rab11 RNA interference in HeLa and HeLa-Rab14-KD2 cells was performed by using QIAGEN HP Custom siRNAs (Rab11a and Rab11b). For LatrunculinA treatment, HeLa-Rab14-KO cells were treated with LatrunculinA overnight at a final concentrations of 1 nM and 4 nM. Stable ACF7 knockdown HeLa cell lines (HeLa-MACF2-KD1 and HeLa-MACF2-KD2) were generated by using Sigma lentiviral human MACF2 shRNA plasmids (TRCN0000300045 and TRCN0000303930). The population was selected with puromycin, and stable clones were isolated.

### Antibodies and plasmids

The following antibodies used for immunofluorescence and western blots: acetylated-α-tubulin (Cell Signalling, D20G3, IF 1:200, WB 1:1000), Rab14 (Sigma-Aldrich, R0656, IF 1:100, WB 1:500), FLAG (Sigma-Aldrich, F1804, IF 1:100, WB 1:1000) and MKLP1 (Thermo Fisher, PA5-31773, IF 1:200). The Alexa-594 and Alexa-647-conjugated anti-mouse and anti-rabbit secondary antibodies used for immunofluorescence were purchased from Life Technologies. The IRDye 680RD Donkey anti-mouse and IRDye 800CW donkey anti-rabbit secondary antibodies used for western blotting were purchased from Li-COR (Lincoln, NE). Phalloidin conjugated to Alexa Flour 594 (A12381) was purchased from Thermo Fisher Scientific (Eugene, Oregon) and LifeAct-mCherry (54491) was bought from Addgene (Michael Davidson). During co-immunoprecipitation, FLAG antibody (Sigma-Aldrich, F3165) was cross-linked to Protein G Sepherose beads. HaloTag Alexa Flour 488 Ligand (G1002) was purchased from Promega (Madison, WI, USA).

### RNA isolation and quantitative PCR

RNA was isolated by using PureLink RNA Mini Kit as per the manufacturers` instructions (Invitrogen, Carlsbad, CA). cDNA was obtained by using the High Capacity RNA to cDNA Kit as per the manufacturers` instructions (Applied Biosystems, Foster City, CA). Quantitative PCR (qPCR) was performed by using TaqMan probes (Applied Biosystems, Pleasanton, CA). Taqman probes used were: Hs03929097 (*GAPDH*) and Hs00156137 (*MACF2*). For quantification, Ct values were normalized to GAPDH and then normalized to control cells.

### Multi-nucleation assay

Control or various KD or KO cells were seeded on collagen I coated (at final 50 μg/ml concentration, Thermo Fisher Scientific, Grand Island, NY) glass coverslips. After 48 hours in culture the cells were fixed in 4% paraformaldehyde (PFA), permeabilized with 0.2% Triton X-100 for 3 min at room temperature (RT) and stained with Alexa Flour 594 phalloidin (Thermo Fisher Scientific, Eugene, Oregon) for 30 min at 37 °C. The coverslips with cells were mounted on 3 cm glass bottom cell culture plates using Vectashield mounting medium with DAPI (Vector Laboratories, Burlingame, CA). Random fields on the coverslips were photographed using an inverted Olympus IX81 microscope (Olympus Europe Holding Gmbh, Germany, Hamburg) with a × 20 oil immersion lens and the number of multi-nucleated cells was counted manually. The rate of total multi-nucleated cells was then calculated. All data is derived from at least three independent experiments.

To rescue multi-nucleation, either HeLa-Rab14-KD2 or HeLa-Rab14-KO cells were seeded on collagen-coated glass coverslips and transfected with various GFP-tagged Rab plasmids. Then 48 hours post-transfection cells were processed and examined for multi-nucleation as described above.

To examine LatrunculinA effect on rescuing multi-nucleation, HeLa-Rab14-KO cells were seeded on collagen-coated glass coverslips and treated with 4 nM concentration of LatrunculinA overnight. After 24 hours cells were processed and examined for multi-nucleation as described above.

### Immunofluorescence and time-lapse microscopy

All fixed cells were imaged with an inverted Olympus IX81 microscope (Olympus Europe Holding Gmbh, Germany, Hamburg) using PlanApo N 60x/1.42 oil lens and the Orca-R^2^ digital camera with excitation system MT10. Image processing was carried out by using the XCELLENCE software.

To measure overall cell division time and the time that cells spend in each mitotic stage, 40,000 cells were plated on fibronectin-coated (at final 30 μg/ml, Sigma Aldrich, CO) 3 cm glass bottom cell culture plates. After 48 hours in culture, the plates were placed in a heat and humidity (37 °C, 5% CO_2_) controlled incubator (INUBG2EONICS Tokai Hit, Shizuoka-ken, Japan) mounted on top of the motorized XY stage. Cells were imaged using an inverted Olympus IX81 microscope (Olympus Europe Holding Gmbh, Germany, Hamburg) with a × 20 oil immersion objective and images were acquired every 10 min. In all cases, the data is derived from at least three independent experiments.

To rescue overall cell division time and the time that cells require to complete each stage of the mitotic phase, either 40,000 control or HeLa-Rab14-KO cells were plated on fibronectin-coated 3 cm glass bottom cell culture plates and transfected with GFP-Rab14 or GFP-Rab11a plasmids. Then 24 hours post-transfection, the cells were analysed by time-lapse imaging as described above.

To determine the effect of Rab11 siRNA on overall cell division time, either control or HeLa-Rab14-KD2 cells were transfected with Rab11a/b siRNAs. Then 24 hours post-transfection cells were plated on fibronectin from bovine plasma (at final 30 μg/ml, Sigma Aldrich, CO) coated 3 cm glass bottom cell culture plates. After 48 hours in culture the cells were analyzed by time-lapse imaging as described previously.

To study CAMSAP3 and Fip3 localization during cell division using time-lapse microscopy, cells were plated on fibronectin-coated glass bottom cell culture plates. After 24 hours in culture the cells were transfected with either GFP-CAMSAP3 or GFP-Fip3 plasmid. Then 24 hours post-transfection, the cells were analyzed by time-lapse imaging as described above.

### Immunoprecipitation and proteomic analysis

Co-immunoprecipitation was performed using HeLa-FLAG-Rab14 and HeLa-FLAG-Rab14-Q70L cell lines. Cells were synchronized using thymidine/nocodazole block as described previously (Schiel *et al*, 2012). Cell were collected at telophase and Triton X-100 lysates were used for immunoprecipitations using anti-FLAG antibody conjugated to protein G sepharose beads and analysed by mass spectrometry as described previously (Schiel *et al*, 2012). All proteins identified by mass spectrometry were filtered using following criteria: (a) not present in IgG control sample and (b) enriched at least 2X in Rab14-Q70L sample. Additionally, we eliminated all RNA and DNA binding proteins, as well as all mitochondria proteins as putative contaminants. The final list is shown in Figure 5A.

### CRISPR/Cas9 mediated GFP-tagging of endogenous Rab14

A HeLa cell line with GFP-tagged endogenous Rab14 was generated via co-electroporation of CRISPR/Cas9 ribonucleoproteins (RNPs) and linear double-stranded DNA homology-directed repair template (HDRT) (see Supplemental Figure 2A), based on the method of Roth et al. (Roth *et al*,). A guide RNA (gRNA) sequence of GTATGGTGCAGTTGCCATGG, which targets in the start codon of Rab14, was selected using the CRISPOR gRNA design tool (Haeussler *et al*,) to identify the gRNA with the highest specificity and efficiency within ~30 nt of the 5’ end of Rab14 exon 1. To generate this gRNA, the crRNA and tracrRNA components were synthesized by Thermo Fisher Scientific (Waltham, MA) and were annealed according to the manufacturer’s protocol. The HDRT consisted of a superfolder GFP (Pedelacq *et al*, ) coding sequence flanked by 350-400 bp homology arms for the genomic regions upstream (the 5’UTR of Rab14) and downstream (Rab14 exon 1 in-frame with GFP, and a portion of intron sequence). This HDRT was synthesized by Integrated DNA Technologies (Coralville, IA), amplified by PCR, and gel-purified. Electroporation was performed using the Neon Transfection System (#MPK5000) and Cas9 protein (#A36497) purchased from Thermo Fisher Scientific. Approximately 5✕10^5^ HeLa cells were electroporated with 25 pmol of RNPs (1:1 Cas9:gRNA ratio) and 2 µg of HDRT using the Neon Transfection System 100 µL Kit (#MPK10096) with pulse conditions of 1150 V/20 ms/2 pulses. The cells were then expanded in culture, and GFP+ cells were isolated from the mixed population at 8 days after electroporation by fluorescence-activated cell sorting.

### Statistical analysis

All statistical analysis was performed using SigmaPlot software (wpcubed GmBH, Germany). For multi-nucleation assay and time-lapse imaging, at least three randomly chosen image fields from a single coverslip were used for data collection. In all cases, data shown are the mean ± s.d. derived from at least three independent experiments. In all figures n represents total number of cells analysed.

## ACKNOWLEDGEMENTS

We would like to thank Dr. Anna Akhmanova (Utrecht University) for providing us with GFP-CAMSAP3 construct, as well as Dr. Jagath Junutula for Rab14 dominant-negative and constitutively-active constructs. This work was supported by NIDDK grant DK064380 to RP and NIAID grant 1R01AI138625 to SVE.

## AUTHOR CONTRIBUTIONS

RP and VAS designed experiments and oversaw entire project. PG performed most of the time-lapse analyses as well as functional assays. PG also generated Rab14 KO and KD lines as well as MACF2 KD lines. EP performed MB proteomic analysis and performed some of the immunofluorescence analyses. HKH and SVE generated GFP-tagged endogenous Rab14 cell line.

---

**Figure S1. Generation of Rab14 knockdown and knockout HeLa cell lines.**

(A-B) Western blots of lysates derived from control, Rab14-KD (A) or Rab14-KO (B) cell lines analysed using anti-tubulin and anti-Rab14 antibodies.

(C) Sequence analysis of two different Rab14-KO cell lines.

**Figure S2. Rab14 localization analysis.**

(A) HeLa cells expressing GFP-tagged endogenous Rab14 were fixed and stained with anti-EEA1 antibody.

(B) GFP tagging of endogenous Rab14 via CRISPR/Cas9 and homology-directed repair (HDR). HeLa cells were electroporated with CRISPR/Cas9 ribonucleoprotein (RNP) with guide RNA (gRNA) targeting the 5’ end of the Rab14 gene, and a homology-directed repair (HDR) template containing a superfolder GFP coding sequence flanked by homology regions (HR) for the target site. Cas9 generates a double-strand break (DSB) at the target site, allowing the template to be integrated at that site by HDR. This results in GFP-Rab14 “knock-in”, by which GFP-tagged Rab14 is expressed from its native genomic locus, avoiding overexpression of the tagged protein. (C) Control and Rab14-KO HeLa cells were fixed and stained with anti-Fip1 antibody.

**Supplemental Figure 3. Generation of MACF2 knockdown HeLa cell lines.**

(A) qPCR analysis of MACF2 mRNA levels in control, MACF2-KD1 and MACF2-KD2 cells.

**Supplemental Figure 4. Uncropped western blots used in this study.**

(A-B) Western blots used in figure 5B.

**Movie 1. Time-lapse analysis of dividing control HeLa cell. Movie 2. Time-lapse analysis of dividing Rab14-KO HeLa cell.**

Time-lapse images shown cell that fails division and generates bi-nuclear cell.

**Movie 3. Time-lapse analysis of dividing Rab14-KO HeLa cell.**

Time-lapse images shown cell with longer time needed to complete cytokinesis.

**Movie 4. Time-lapse analysis of dividing control HeLa cell expressing GFP-FIP3.**

Movie shows merged bright-field and GFP-FIP3 channels.

**Movie 5. Time-lapse analysis of dividing control HeLa cell expressing GFP-FIP3.**

Movie shows only GFP-FIP3 channel. For merged channels see supplemental movie 4.

**Movie 6. Time-lapse analysis of dividing Rab14-KO1 HeLa cell expressing GFP-FIP3.**

Movie shows merged bright-field and GFP-FIP3 channels.

**Movie 7. Time-lapse analysis of dividing Rab14-KO1 HeLa cell expressing GFP-FIP3.**

Movie shows only GFP-FIP3 channel. For merged channels see supplemental movie 6.

**Movie 8. Time-lapse analysis of dividing MACF2-KD1 HeLa cell.**

Movie shows cell with longer time needed to complete cytokinesis.

**Movie 9. Time-lapse analysis of dividing MACF2-KD1 HeLa cell expressing GFP-FIP3.**

Movie shows merged bright-field and GFP-FIP3 channels.

